# Single cell transcriptome analysis reveals RGS1 as a new marker and promoting factor for T cell exhaustion in multiple cancers

**DOI:** 10.1101/2019.12.26.888503

**Authors:** Yunmeng Bai, Meiling Hu, Zixi Chen, Jinfen Wei, Hongli Du

## Abstract

T cell exhaustion is one of the main reasons of tumor immune escape. Using single cell transcriptome data of CD8+ T cells in multiple cancers, we identified different cell types, in which Pre_exhaust and exhausted T cells participated in negative regulation of immune system process. By analyzing the co-expression network patterns and differentially expressed genes of Pre_exhaust, exhausted and effector T cells, we identified 35 genes related to T cell exhaustion, which high GSVA scores were associated with significantly poor prognosis in various cancers. In the differentially expressed genes, *RGS1* showed the greatest fold change in Pre_exhaust and exhausted cells of three cancers compared with effector T cells, and high expression of *RGS1* was also associated with poor prognosis in various cancers. Additionally, RGS1 protein was upregulated significantly in tumor tissues in the immunohistochemistry verification. Furthermore, *RGS1* displayed positive correlation with the 35 genes, especially highly correlated with *PDCD1, CTLA4, HAVCR2* and *TNFRSF9* in CD8+ T cells and cancer tissues, indicating important roles of *RGS1* in CD8+ T cell exhaustion. Considering the GTP-hydrolysis activity of *RGS1* and significantly high mRNA and protein expression in cancer tissues, we speculated that *RGS1* potentially mediate the T cell retention to lead to the persistent antigen stimulation, resulting in T cell exhaustion. In conclusion, our findings suggest that RGS1 is a new marker and promoting factor for CD8+ T cell exhaustion, and provide theoretical basis for research and immunotherapy of exhausted cells.

## 1. Introduction

T cell exhaustion (Tex), a hyporesponsive state of T cells with increased inhibitory receptors, decreased effector cytokines and impaired cytotoxicity, was originally described in CD8+ T cells during chronic lymphocytic choriomeningitis virus (LCMV) of mice(1). In recent years, the phenomenon of Tex has also been found in cancers(2, 3), which is one of the main reasons of tumor immune escape(4). It has been reported that exhausted T cells in cancers share many similarities with that in chronic infection(5), and play a significant role in tumorigenesis(6). Studies show that exhausted T cells can be used as one of the main targets of immunosuppression therapy to save T cell from exhaustion and reactivate the cytotoxicity of T cells, providing a new opportunity for clinical immunotherapy(7). Nevertheless, due to the complexity and heterogeneity of cancers, the concrete mechanisms and molecules of T cell exhaustion in cancers have not been fully elucidated.

Currently, single-cell RNA sequencing (scRNA-seq) has clearly revealed some new mechanisms and phenomena of cancer with the advantages of high accuracy and reproducibility(8–10). Using single cell transcriptome profiling, we can identify new types of immune cells which can’t be revealed at the original tissue level and can construct a developmental trajectory for immune cells which can reveal the heterogeneity(11). These new findings are useful to better understand the immune system and its mechanism of action on tumors. Notably, this technology makes it possible to explore complicated tumor microenvironment including tumor-infiltrating lymphocytes (TILs) in melanoma, head and neck cancer, breast cancer and glioblastoma cancer(12–15). Thus, using advantage of scRNA-seq to analyze T cells and obtain the hallmarks of exhausted T cells can bring a new therapeutic strategy on clinical cancer treatment.

Due to the vital role of CD8+ T cells in eliciting antitumor responses(16), we integrated single cell transcriptome data from colorectal cancer (CRC), hepatocellular cancer (HCC), and non-small cell lung cancer (NSCLC) to analyze CD8+ T cells in various cancers in the present study. Focusing on CD8+ T cell exhaustion associated clusters, we identified *RGS1* as a new marker and promoting factor for T cell exhaustion in multiple cancers with poor prognosis, and showed highly positive correlation with the well-known genes associated with T cell exhaustion. Besides, RGS1 protein highly expressed in tumor tissues was also verified in the immunohistochemistry (IHC) experiment. Our findings could facilitate to understand the mechanism during formation and development of T cell exhaustion, and provide theoretical basis for research and immunotherapy of exhausted cells.

## 2. Materials and Methods

### 2.1 Data acquisition

The single-cell gene expression matrices including raw count and TPM data were obtained from the GEO database: GSE108989 (CRC), GSE99254(HCC), GSE98638(NSCLC), and we isolated CD8+ T cells from peripheral blood (P), adjacent normal (N) and tumor tissues (T).

### 2.2 Quality control and data processing

The raw count expression matrices were processed by R package Seurat v3.2.0 (http://satijalab.org/seurat/). To filtered out the low-quality cells, we excluded cells with < 600 and > 10000 detected genes(17). Counts were log-normalized, and scaled by linear regression against the number of reads with function NormalizeData and ScaleData. The highly variable genes (HVGs) were generated with FindVariableFeatures. Principal component analysis (PCA) was performed on the top 2000 HVGs using functions RunPCA. The appropriate PCs were selected for graph-based clustering with function FindNeighbors and FindClusters. For visualization of clustering analysis, we performed uniform manifold approximation and projection for dimension reduction (UMAP) using RunUMAP function in Seurat. To eliminate the obvious effect from different patients, we performed standard normalization and variable feature selection after acquiring the data. Next, the function FindIntegrationAnchors was performed to find a set of anchors, which used to integrate the data by the function IntegrateData. Besides, the function AddModuleScore was used to calculate scores of gene list in different cell types.

### 2.3 Cell type annotation

Differentially expressed genes (DEGs) of each cluster were identified based on Wilcoxon rank-sum test using function FindAllMarkers compared with the rest clusters. In brief, for each cluster, only genes that met these criteria were considered as cluster-specific DEGs: 1) log2FC > 0.25; 2) expressed > 25% in either of the two groups of cells; 3) adjusted p value < 0.05. The top DEGs were selected to annotated of each cluster based on the canonical markers from previous studies, besides, the CellMarker databases(18) and R package SingleR v1.4.0(19) were performed to further improve the accuracy of the annotation.

### 2.4 Trajectory Analysis

To explore the potential functional changes of CD8+ T cell of different clusters for each cancer, we performed development trajectory analysis by R package Monocle v2.18.0(20) with the cluster-specific genes of each cluster. Dimensional reduction and cell ordering were performed using reduceDimension and orderCells function with default parameter.

### 2.5 Gene ontology (GO) enrichment analysis of DEGs

Biological significance was explored by GO term enrichment analysis by R package clusterProfiler v3.18.0(21), biological process, cellular component, and molecular function included. Adjusted p value < 0.05 was considered statistically significant. Visualization is realized by R package ggplot2 v3.3.3 and ggalluvial v0.12.3(22).

### 2.6 Weighted gene co-expression network analysis

In order to identify the highly linked genes in specific clusters, weighted gene co-expression network analysis (WGCNA) was performed with functions in the R package WGCNA(23). To attenuate the effects of noise and outliers, we constructed pseudocells(24) which calculated as averages of 10 cells randomly chosen within each cluster. The function pickSoftThreshold was used to calculate the soft power parameter and blockwiseModules to construct co-expression network. Finally, corPvalueStudent was used to mine modules related to specific cell types.

### 2.7 Survival analysis

According to the median of gene expression values, cells were divided into high and low groups, then the survival curve was shown using the Kaplan–Meier curve with a log-rank test by GEPIA2 (http://gepia2.cancer-pku.cn/) to illustrate the relationship between differential genes and overall patient survival. The terms with P-value < 0.05 were identified as significant. Additionally, multiple hypothesis testing (FDR) was performed to the significant p value using the function p.adjust.

### 2.8 Gene set variation analysis

For generated gene set, we performed gene set variation analysis (GSVA) to the data in the present study and validation dataset using R package GSVA(25) with the default parameter, to evaluate the effect distinguishing Tex cells.

### 2.9 The basic expression of mRNA and protein in normal and cancer tissues

The RNA-seq data were downloaded from TCGA database (https://portal.gdc.cancer.gov/) and then calculated into TPM value. According to the annotation, samples were grouped into normal and four stages of cancer groups. The differential expression of mRNA in the different groups was performed by wilcoxon test with p value<0.05.

The protein expression level in normal and cancer tissues was analyzed using Human Protein Atlas (HPA) database (https://www.proteinatlas.org). Immunohistochemistry pictures were downloaded from the Tissue Atlas and Pathology Atlas.

### 2.10 Immunohistochemistry

Immunohistochemistry was carried out on human liver cancer tissue microarray (Shanghai Outdo Biotech Co., Ltd, China). The tissue sections were first dried at 63□ for 1 h, dewaxed and rehydrated before epitope-retrieval by heating at 100 C in 10 mM sodium-citrate (pH 6.0) for 5 min in EDTA solution for 20 min. The sections were cooled down to room temperature for 30 min. The tissue sections were treated with 3% hydrogen peroxide for 20 min to eliminate the endogenous peroxidase and alkaline phosphatase activity in the tissue. After cooling down to room temperature, the sections were treated by blocking agents for 10 min. The sections were incubated with of the individual primary antibody (RGS1, Invitrogen, USA; Product #PA5-86730,) diluted 1:1000 800 overnight at 4□, followed by secondary antibodies at room temperature for 30 min. DAB(3,3’-diaminobenzidine) was then applied as a substrate to reveal the antigen. Hematoxylin was used for counterstaining. The stained images were counted by Image J software(26), and the optical density (OD) value was used for quantification.

## 3. Results

### 3.1 Single CD8+ T cell transcriptome landscape

We obtained single cell transcriptome data of human T cells from GEO database, including 12 patients from CRC(27), 6 patients from HCC(28) and 14 patients from NSCLC(29). After strict quality control and filtration, we collected 4010, 1752 and 4439 CD8+ T cells from peripheral blood (P), adjacent normal (N) and tumor tissues (T) (Supplemental Figure 1A, 1B, Supplemental Table 1).

We divided the cells of each cancer into different cell types annotated with cluster-specific genes expression (Figure 1A, Supplemental Table 2). Specifically, in CRC, cells were identified as naïve T cells (Tn, cluster 4, marked with selectin L (*SELL*), lymphoid enhancer binding factor 1(*LEF1*), C-C motif chemokine receptor 7(*CCR7*)), transcription Factor 7 (*TCF7*), effector T cells (Teff, cluster 2 and cluster 5, marked with C-X3-C motif chemokine receptor 1(*CX3CR1*), killer cell lectin like receptor F1(*KLRF1*)), fibroblast growth factor binding protein 2 (*FGFBP2*), Fc fragment of IgG receptor IIIa (*FCGR3A*), exhausted T cells (Tex, cluster 3 and cluster 6, marked with hepatitis A virus cellular receptor 2(*HAVCR2*), programmed cell death 1(*PDCD1*), lymphocyte activating 3 (*LAG3*), *TOX, CXCL13*, tissue-resident memory T cells (Trm, cluster 1, marked with *CD69*, integrin subunit alpha E (*ITGAE*)), mucosal-associated invariant T cells (MAIT, cluster 7, marked with solute carrier family 4 member 10(*SLC4A10*)), RAR related orphan receptor C (*RORC*) (Figure 1B, 1C). Cluster 0 was located between cluster 2 and cluster 3, which represented as Teff and Tex cells, besides, the genes marked Teff and Tex cells showed relative high expression levels in this cluster, therefore, we identified it as Pre_exhaust T cells. Similarly, cells in HCC were identified as Tn (cluster 4), Teff (cluster 0), Pre_exhaust (cluster 1), Tex (cluster 3 and cluster5), MAIT (cluster 2), cells in NSCLC were identified as Tn (cluster 4), Teff (cluster 0 and cluster 6), Pre_exhaust (cluster 1 and cluster 7), Tex (cluster 3), Trm (cluster 2), MAIT (cluster 5). Different cell types showed preference in different tissues. In general, Tn and Teff cells were mainly enriched in peripheral blood, Pre_exhaust and Tex cells were mainly enriched in tumor tissues, Trm cells existed more in adjacent normal tissues of CRC and HCC, and in tumor tissues of NSCLC, which may be related to tissue specificity. Besides, the number of MAIT cells was relative less than the others. Additionally, we performed exhausted cell scoring by the function AddModuleScore in R package Seurat using the exhaustion gene list including *HAVCR2*, T cell immunoreceptor with Ig and ITIM domains (*TIGIT*), *LAG3, PDCD1, CXCL13*, layilin (*LAYN*), *TOX*, cytotoxic T-lymphocyte associated protein 4 (*CTLA4*), B and T lymphocyte associated (*BTLA*) and visualized in Supplemental Figure 1C, showing the accuracy of Tex cell classification (Supplemental Figure 1C).

**Figure 1.**
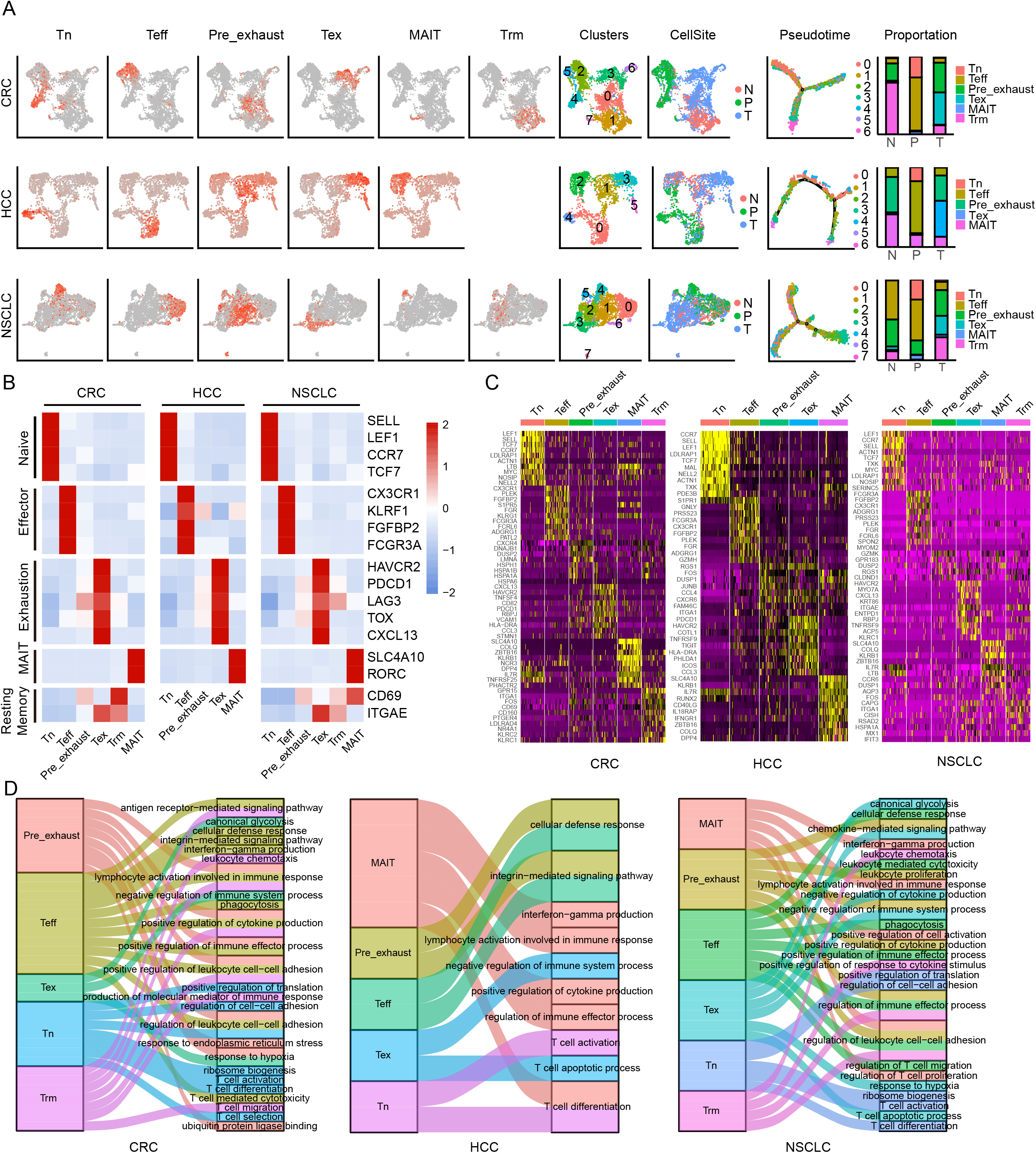
Clustering of CD8+ T cells in three cancers. **A**. UMAP of single cells to visualize cell-type clusters (left), Pseudo-time trajectory graph (middle) and the proportion of different cell types in different sources (right). **B**. Heatmap showing marker genes for CD8+ cell types. **C**. The top 10 DEGs in each cell type in three cancers. **D**. The GO enrichment analysis of different cell types of CD8+ T cells in three cancers.

To further explore the function of each cluster, we performed GO enrichment analysis using the cluster-specific genes (Figure 1D). In all three cancers, Tn cells were enriched in T cell activation, T cell differentiation, ribosome biogenesis; Teff cells were enriched in cellular defense response, positive regulation of cytokine production, T cell mediated cytotoxicity; Tex cells were enriched in negative regulation of immune system process, T cell apoptotic process, response to hypoxia; Trm cells were enriched in antigen receptor-mediated signaling pathway, leukocyte chemotaxis; MAIT cells were enriched in interferon-gamma production; and Pre_exhaust cells were enriched in terms associated with Teff and Tex cells, including cellular defense response and negative regulation of immune system process, showing the transitional characteristics during T cell exhaustion.

### 3.2 Establishment of co-expression network

Tex cells, as shown above, played a negative role in immune system process, and it was suggested that Pre_exhaust cells was a transitional stage from Teff to Tex cells. To find out the highly linked genes associated with T cell exhaustion, we used the R package WGCNA to construct the weighted co-expression network in Teff, Pre_exhaust and Tex cells (Figure 2A). The blue module (R-value 0.96, P-value 2e-145), turquoise module (R-value 0.96, P-value 7e-67) and blue module (R-value 0.96, P-value 2e-172) represented Tex cells in CRC, HCC and NSCLC, receptively. Combining with the differentially expressed genes (DEGs) up-regulated in the Tex vs. Teff cells comparison in three cancers (Supplemental Table 3), there were 35 genes were found in three Tex cells-related modules and overexpressed in Tex cells (Figure 2B, Table 1), which were defined as “Candidate” gene set, including exhaustion marker *PDCD1, CTLA4, HAVCR2, TOX, TIGIT*. It is noteworthy that the housekeeping gene glyceraldehyde-3-phosphate dehydrogenase (*GAPDH*), which can catalyze an important energy-yielding step in glycolysis metabolism, was also upregulated and enriched in the highly linked DEGs, thus we speculated that the glycolysis progress was disordered in Tex cells.

**Figure 2.**
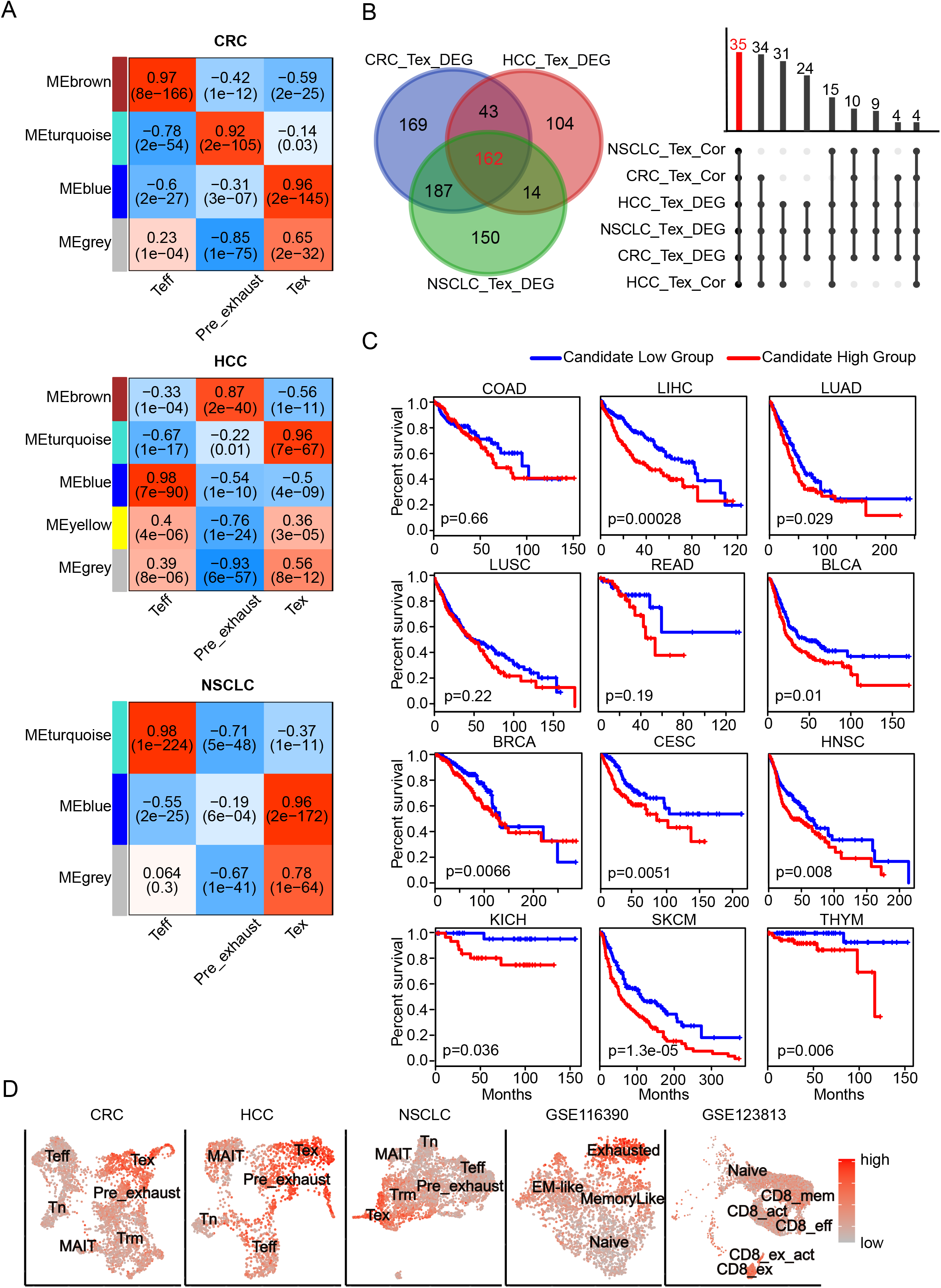
Candidate gene set associated with CD8+ Tex cells. **A**. The gene co-expression network modules of CD8+ T cells with correlation coefficient and P-value. **B**. The number of genes with differential expression (left) and co-expression (right). **C**. The Kaplan-Meier overall survival curves of TCGA patients grouped by the middle expression value of Candidate gene set. The red and blue lines denote higher and lower expression group, respectively. **D**. Distinguishing Tex cells from the other CD8+ T cells effectly in different cancers by GSVA score of Candidate gene set.

**Table 1.**
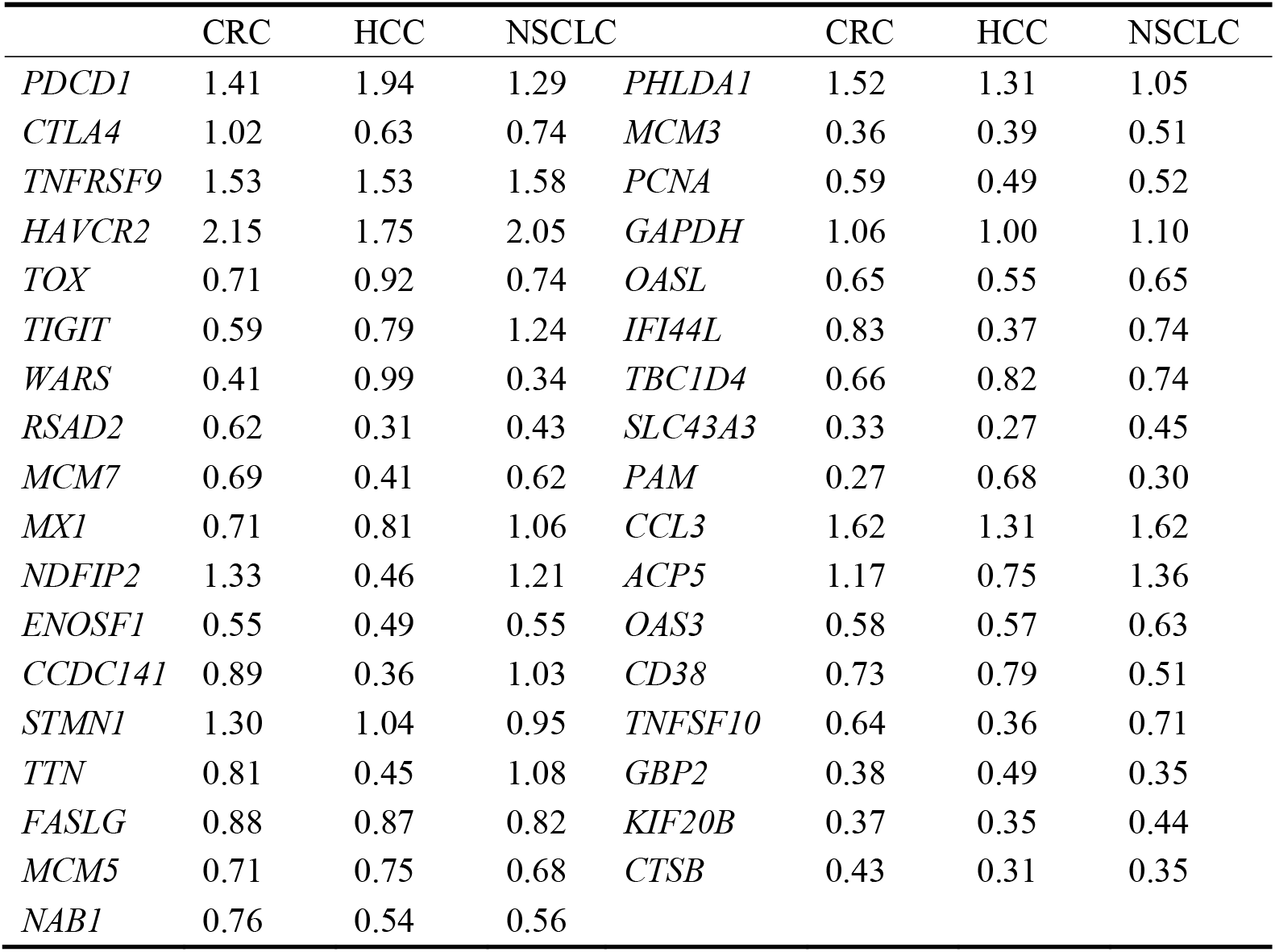
The logarithm of fold change of Candidate gene set in three cancers.

As shown in Figure 2C, poor overall survival was correlated with higher Candidate gene set expression in multiple cancers including liver hepatocellular carcinoma (LIHC), lung adenocarcinoma (LUAD), bladder urothelial carcinoma (BLCA), breast invasive carcinoma (BRCA), cervical squamous cell carcinoma and endocervical adenocarcinoma (CESC), head and neck squamous cell carcinoma (HNSC), kidney chromophobe (KICH), skin cutaneous melanoma (SKCM), thymoma (THYM). Besides, the GSVA scores of Candidate gene set in Tex cells were much higher than those of the other clusters in CRC, HCC and NSCLC, so were in the other independent validation datasets (Figure 2D). GSE116390 dataset(30) was composed of four types of tumor-infiltrating CD8+ T cells, which were exhausted, memory-like, naïve, and effector memory-like (EM-like) subsets, from B16 melanoma tumor-bearing mice, and GSE123813 dataset(31) contained six types of tumor-infiltrating CD8+ T cells, which were from 11 patients with advanced basal cell carcinoma, showing higher score in the exhausted related clusters.

As indicated above, the Candidate gene set was enriched in exhausted CD8+ T cells with poor prognosis and was able to distinguishing Tex cells from the other CD8+ T cells in different cancers, indicating the GSVA score of these 35 genes might be an effective prognostic marker or a marker to identify Tex cells.

### 3.3 Differentially expressed genes associated with T cell exhaustion

The exhaustion of T cells was gradually formed, so we performed differential expression analysis of Pre_exhaust and Tex cells compared with Teff cells, in order to further explore the critical genes associated with the formation and development of T cell exhaustion. In the Pre_exhaust vs. Teff cells comparison, there were 119 DEGs existed in all three cancers, 57 of whom were up-regulated and the others were down-regulated. Correspondingly, there were 162 up-regulated and 88 down-regulated DEGs in the Tex vs. Teff cells comparison. Furthermore, there were 40 DEGs overexpressed in both Pre_exhaust and Tex cells compared with Teff cells (Figure 3A), which may contribute to the origin of T cell exhaustion, including the canonical exhaustion marker *PDCD1* which encodes an inhibitory receptor(32), *CD69* with the capability to mediate the cell retention(33), Cbl Proto-Oncogene B(*CBLB*) whose deletion can inhibit CD8+ T cell exhaustion(34), hypoxia inducible factor-1 (*HIF1A*) which can stably expressed in hypoxia condition, consistent with the biological process of Tex cells (Figure 1D).

**Figure 3.**
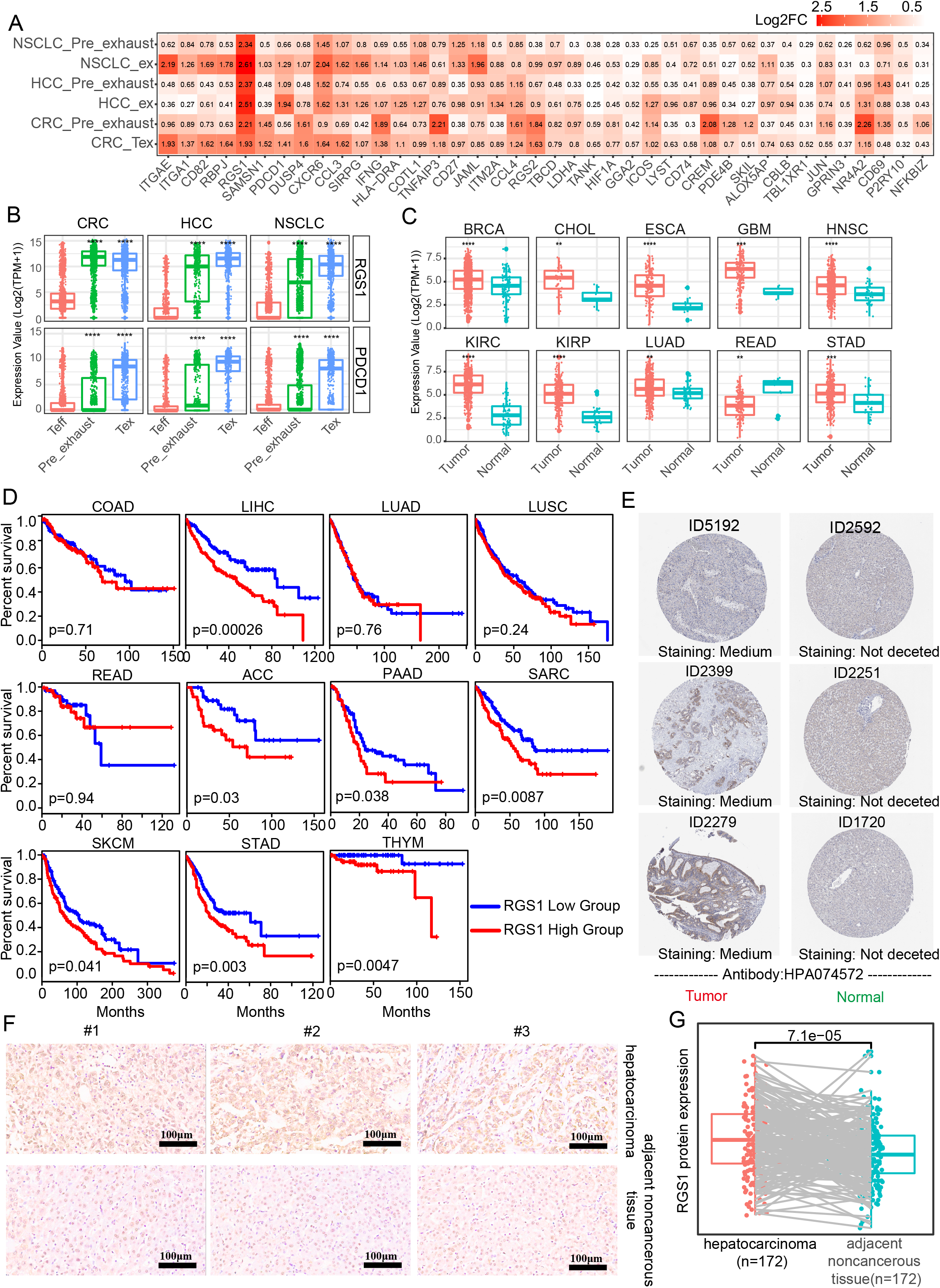
DEGs in Pre_exhasuted and Tex cells compared with Teff cells. **A.** The common up-regulated genes in Pre_exhasuted and Tex cells compared with Teff cells. **B**. The mRNA expression value of *RGS1* in single cell dataset and TCGA database (**C**). **D**. The Kaplan-Meier overall survival curves of TCGA patients grouped by the middle expression value of *RGS1.* The red and blue lines denote higher and lower expression group, respectively. **E**. Representative IHC images of RGS1 protein in tumor and normal tissues of liver derived from the HPA database and verification experiment (**F**, scale bar 100 μm, magnification ×20). **G**. The protein expression value of RGS1 in hepatocarcinoma and adjacent noncancerous tissues in the IHC verification experiment.

Obviously, regulator of G protein signaling 1(*RGS1*) showed the greatest fold change in Tex cells across three cancers and also across different patients, showed the greatest fold change in Pre_exhaust cell in HCC and NSCLC (Figure 3A, Supplemental Figure 2), illustrating its potential roles during T cell exhaustion. Besides, *RGS1* was expressed highly in whole Tex cells compared with Teff cells (Figure 3B), eliminating that the high fold change of *RGS1* in Pre_exhaust and Tex cells was not caused by partial cells with abnormally high expression value but the high expression in whole cells. In order to evaluate the role of *RGS1* in tumorigenesis, we analyzed the expression levels of *RGS1* between tumor and normal tissues in TCGA database (Figure 3C). *RGS1* expression was significantly upregulated in multiple cancers including BRCA, cholangiocarcinoma (CHOL), esophageal carcinoma (ESCA), glioblastoma multiforme (GBM), HNSC, kidney renal clear cell carcinoma (KIRC), kidney renal papillary cell carcinoma (KIRP), LUAD, rectal carcinoma (READ) and stomach adenocarcinoma (STAD) when compared with the normal samples. Besides, we added and compared the expression levels of *RGS1* across different cancer stages, and found *RGS1* expression was associated with stage in some cancer types, such as, *RGS1* expression was higher in Stage II, III and IV vs Stage I in STAD. This result revealed that *RGS1* was likely a key tumorigenesis regulator in multiple cancers and may associated with prognosis. Conspicuously, the prognosis analysis was analyzed in 33 TCGA cancer types (Supplemental Table 4). RGS1 expression was significantly correlated to poor prognosis in seven cancers, including LIHC, adrenocortical carcinoma (ACC), pancreatic adenocarcinoma (PAAD), sarcoma (SARC), SKCM, STAD and THYM (Figure 3D), suggesting that *RGS1* was a potential prognostic factor in survival of above cancers. Apart from that, the protein level of RGS1 in HPA database showed the immunohistochemical (IHC) staining of RGS1 was negative staining in normal tissues and positive in liver cancer tissues, demonstrating that RGS1 was significantly expressed in cancer tissues than in normal liver tissues (Figure 3E). Additionally, we performed IHC verification to quantify the protein expression of RGS1 using the local clinical samples of liver cancer and normal tissues (Figure 3F, 3G), similarly, RGS1 protein displayed stronger staining in hepatocarcinoma, line with the statistical result (p=7.1e-5). The clinicopathological information of the patient samples and protein expression value were provided in Supplemental Table 5. These results showed that *RGS1* was highly expressed in Tex cells in cancers, upregulated in tumor tissues in mRNA and protein level, and with poor prognosis in multiple cancers, which indicated its potential key role in T cells exhaustion or cancer progress, and RGS1 might be an effective prognostic marker or a marker to identify Tex cells.

### 3.4 Correlation between RGS1 and Candidate gene set

To discover the relationship between *RGS1* and Candidate gene set as their potential roles in exhausted T cells, we calculated the correlation coefficient in single cell and tissue level (Figure 4). In three cancer single cell datasets, it was obvious that *RGS1* showed positive correlation with 35 genes (up to 0.3~0.8), indicating that consistency of coexpression patterns between these genes. In the bulk RNA sequencing datasets, the correlation coefficients were almost positive, especially high with *PDCD1, CTLA4*, TNF receptor superfamily member 9 (*TNFRSF9*), *HAVCR2, TOX* and *TIGIT*, which further confirmed the potential key roles of *RGS1* in Tex cells. The negative correlations were mainly with the genes involved in cell cycle and DNA replication, including minichromosome maintenance complex component 7 (*MCM7*), enolase superfamily member 1 (*ENOSF1*), minichromosome maintenance complex component 5 (*MCM5*), minichromosome maintenance complex component 3 (*MCM3*), proliferating cell nuclear antigen (*PCNA*), which pointed out the different results observed between single cell and bulk RNA sequencing analysis, suggesting we could obtain deeper understanding about T cell exhaustion by single cell sequencing. One interpretation of above inconsistent result is that the bulk transcriptome change (which mixes many immune and non-immune cells together) likely reflects an overall expression in these cells and does not discriminate specific T cell states, further highlights the advantages in defining and studying T cell exhaustion state by using single cell sequencing data.

**Figure 4.**
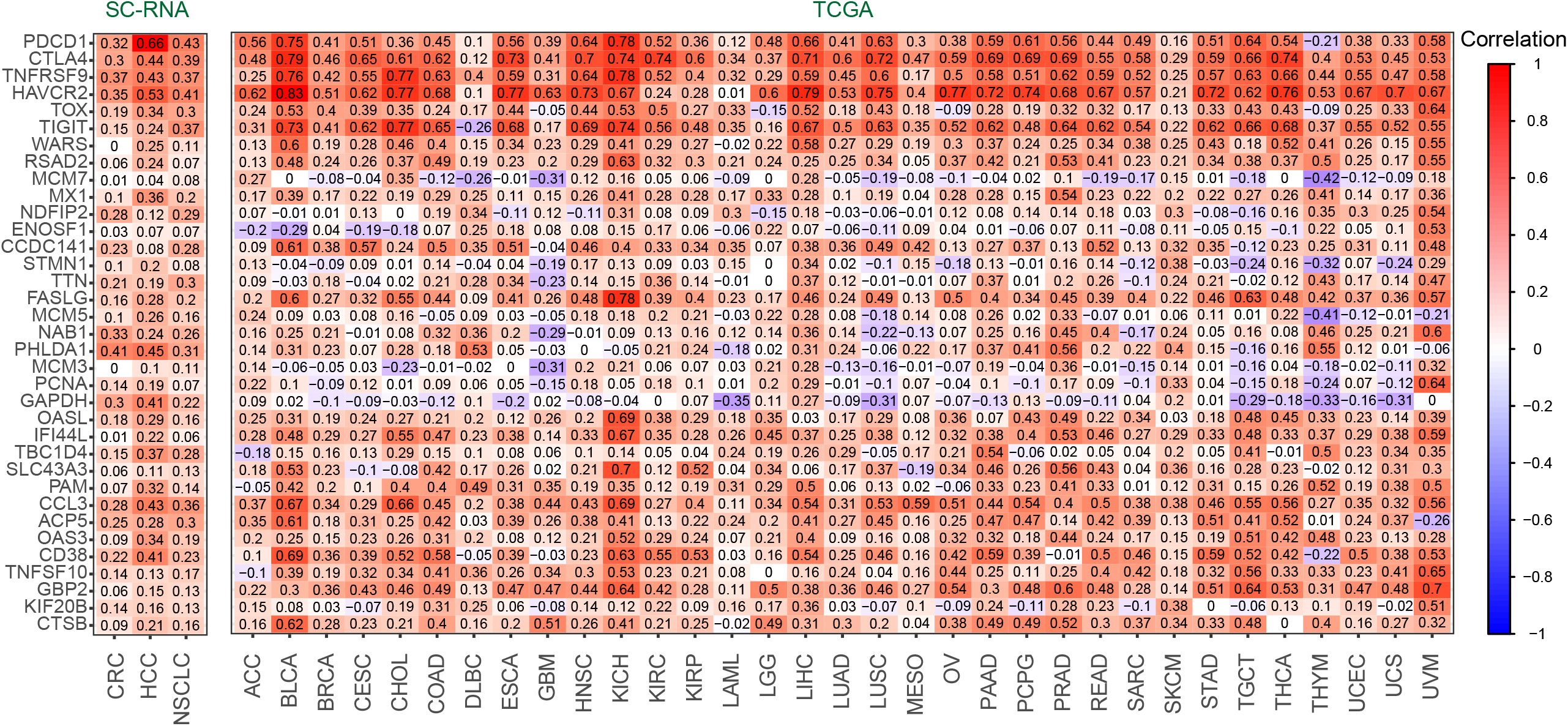
The correlation coefficient between *RGS1* and Candidate gene set of Tex cells in CD8+ T cells (left) and TCGA database(right).

## 4. Discussion

T cell exhaustion is characterized by loss of effector functions, continuously high expression of numerous inhibitory receptors, epigenetic and transcription profile changes, and dysregulated metabolism. Exhaustion CD8+ T cells are associated with suppressive immune microenvironment and poor overall survival in various cancer types, such as, in invasive bladder cancer(35) and clear-cell renal cell carcinoma(36). Besides, increased exhausted CD8+ T cell subpopulations predict PD-1 blockade resistance response in melanoma(37). Accumulating evidences support exhausted T cells are possible to be rescued in cancer immunotherapy. Anti-PD1 antibodies, including atezolizumab and nivolumab, can renew the activity of exhausted CD8 T cells through preventing PD-1-mediated attenuation of proximal TCR cascades(38, 39) and can effect metabolic reprogramming to reinvigorate T cells(40). However, one study found an association between increased accumulation of one CD8+ T cell exhaustion phenotype and clinical benefit, suggesting exhausted T cells may comprise heterogenous cell population with distinct responsiveness to intervention and the standard definition of exhaustion cells is unclear in the context of treatment(41). Thus, understanding molecular mechanism of T cell exhaustion and comprehensively exploring potential markers associated with T cell exhaustion is essential to precisely define T cell exhaustion and establish rational immunotherapeutic interventions.

In this study, combining with single-cell RNA sequencing, which can facilitate to detect he transcriptome on the level of single cell(42), we can shed light on the complication of tumor-infiltrating T cells. In order to explore the key genes associated with T cell exhaustion in multiple cancers, we performed transcriptomic analysis of single CD8+ T cells isolated from three cancers, including CRC, HCC and NSCLC, and identified different cell types, thereinto, Pre_exhaust and Tex cells overexpressed exhaustion markers and enriched in the negative regulation of immune progress. In the comparison with Teff cells, *RGS1* showed almost the greatest fold change in Pre_exhaust and Tex cells of three cancers with poor prognosis, and displayed highly positive correlation with the well-known genes associated with T cell exhaustion.

In the WGCNA analysis, we identified a Candidate gene set consisting of 35 DEGs, including exhaustion markers such as *PDCD1, CTLA4, HAVCR2, TOX, TIGIT*. Apart from that, genes involved in cell cycle and DNA replication also included, such as *MCM7, MCM5, MCM3, PCNA*, stathmin 1 (*STMN1*), titin (*TTN*), TBC1 domain family member 4 (*TBC1D4*), suggesting that exhausted cells still retained the ability of proliferation, which was also observed in chronically infected models(43). Functionally, the Candidate gene set was able to distinguish Tex cells from the other subtypes of CD8+ T cells in different cancers and higher GSVA scores of Candidate gene set showed poor prognosis in multiple cancers.

Besides, several DEGs appeared in Pre_exhaust and Tex cells, suggesting the role in the formation and development of T cell exhaustion, in which *RGS1* showed almost the greatest fold change. *RGS1* encodes a member of the regulator of G-protein signaling family, can act as a GTPase activating protein (GAP), increasing the rate of conversion of the GTP to GDP, driving G-protein into its inactive GDP-bound form, hence attenuating or turning off G-protein-coupled receptors signaling(44). *RGS1* is highly expressed in immune cells including T cells(45), B cells(46), natural killer (NK) cells(47), dendritic cells(48) and monocytes(49), suggesting a role for RGS1 in immune cell regulation. RGS1 inhibits the chemokine-induced lymphocyte migration(50) because chemokine-dependent activation of G-protein-coupled receptors can cause the activation of heterotrimeric G-protein subunits resulting in enhanced cell migration and adhesion(51), which has been found in Treg cells(45). In the present study, *RGS1* was highly expressed in tumor tissues and correlated with shorter overall survival, which also appeared in several previous studies, including multiple myeloma(52), melanoma(53, 54), non-small cell lung cancer(55), gastric cancer(56), diffuse large B cell lymphoma(57) and so on. However, the role of *RGS1* in CD8+T cells especially in Tex cells has not been reported. In addition, RGS1 protein, locating at cytoplasm and membrane, enriched in tumor tissues compared with normal tissues according to the IHC staining in HPA database and verification experiment, further verifying its pathogenicity. Considering the ability to block cell migration of *RGS1*, we speculate that *RGS1* can mediate the cell retention to lead to the persistent antigen stimulation of T cells, which resulted in T cell exhaustion with the overexpression of inhibitory genes such as *PDCD1* and *HAVCR2*(58). Additionally, *RGS1* was identified as a HIF-dependent hypoxia target that dampens cell migration and signal transduction(59), indicating its role in exhausted T cells might be caused by hypoxia condition (Figure 1B).

*RGS1*, the most up-regulated gene in Pre_exhaust and Tex cells and a potential marker for T cell exhaustion, was excluded from the Candidate gene set. This phenomenon happened in other T cell exhaustion-related genes as well, such as *CD69* and *CBLB. CD69*, upregulated in Pre_exhaust and Tex cells, is an early activation marker of T cells(60). It can mediate the cell retention via the interaction with sphingosine-1-phosphate receptor 1 (*S1PR1*) which acts as a central mediator of lymphocyte output(61), leading to the persistent antigen stimulation of T cells, which resulted in T cell exhaustion by overexpression of *PDCD1* and *HAVCR2* (62). *CBLB*, also upregulated in Pre_exhausted and Tex cells, deletion can inhibit CD8+ T cell exhaustion and promote chimeric antigen receptor T-cell function(34). Considering the positive correlation between *RGS1* with Candidate gene set in single cells and tissues and its high expression in Tex cells, it is important and the necessity to further study *RGS1* mechanism in T cell exhaustion.

In summary, our findings suggest that the GSVA score of the 35 Candidate gene set could be an effective prognostic marker or a marker to identify Tex cells. *RGS1*, as the most up-regulated gene in Pre_exhaust and Tex cells, might play key roles in T cells exhaustion or cancer progress. As a HIF-dependent hypoxia target, *RGS1* might be upregulated by hypoxia, and further mediate the cell retention by inhibiting chemokine-induced lymphocyte migration. The current study could provide theoretical basis for research and immunotherapy of exhausted cells, while further studies are essential to fully elucidate the concrete mechanism of *RGS1* during CD8+ T cell exhaustion.

## Author Contributions

All authors contributed significantly to the work and the preparation of the manuscript. HD conceived the study. YB analyzed the data with assistance from ZC, wrote the manuscript. MH collected the data and performed IHC experiment. JW reviewed and revised the manuscript. All authors contributed to the article and approved the submitted version.

## Acknowledgments

This research was funded by the National Key R&D Program of China (2018YFC0910200), the Key R&D Program of Guangdong Province (2019B020226001)

## Data Availability Statement

The original contributions presented in the study are included in the article/Supplementary Material, the original IHC figures were uploaded in https://www.jianguoyun.com/p/DdToBqoQ4ISAChiXhJkE and further inquiries can be directed to the corresponding author/s.

## Supplemental Information titles and legends

**Supplemental Figure 1: Clustering of CD8+ T cells in three cancers.**

**A.** UMAP visualization of T cells in three cancers.

**B.** The number of detected genes in each cell before filter in three cancers.

**C.** The exhaustion scores of different cell types using the exhaustion gene list including *HAVCR2, TIGIT, LAG3, PDCD1, CXCL13, LAYN, TOX, CTLA4, BTLA*.

**Supplemental Figure 2: The correlation coefficient between *RGS1* and Candidate gene set of Tex cells in CD8+ T cells of different patients.**

**Supplemental Figure 3: The mRNA expression value of *RGS1* of different stages in TCGA database.**

**Supplemental Table 1: The patient ID and numbers of cell in three cancers**

**Supplemental Table 2: Differentially expressed genes of each cluster in three cancers**.

**Supplemental Table 3: Differentially expressed genes of Pre_exhaust and Tex cells compared with Teff cells in three cancers**.

**Supplemental Table 4: Survival analysis results of .RGS1 in different cancer types.** Adj.p value referred to the adjusted p value corrected by multiple hypothesis testing (FDR).

**Supplemental Table 5: The clinicopathological information of the patient samples and RGS1 protein expression value.**

