## Supplementary figures and images for "Single cell transcriptome analysis reveals RGS1 as a new marker and promoting factor for T cell exhaustion in multiple cancers"

### Supplemental Figure 2

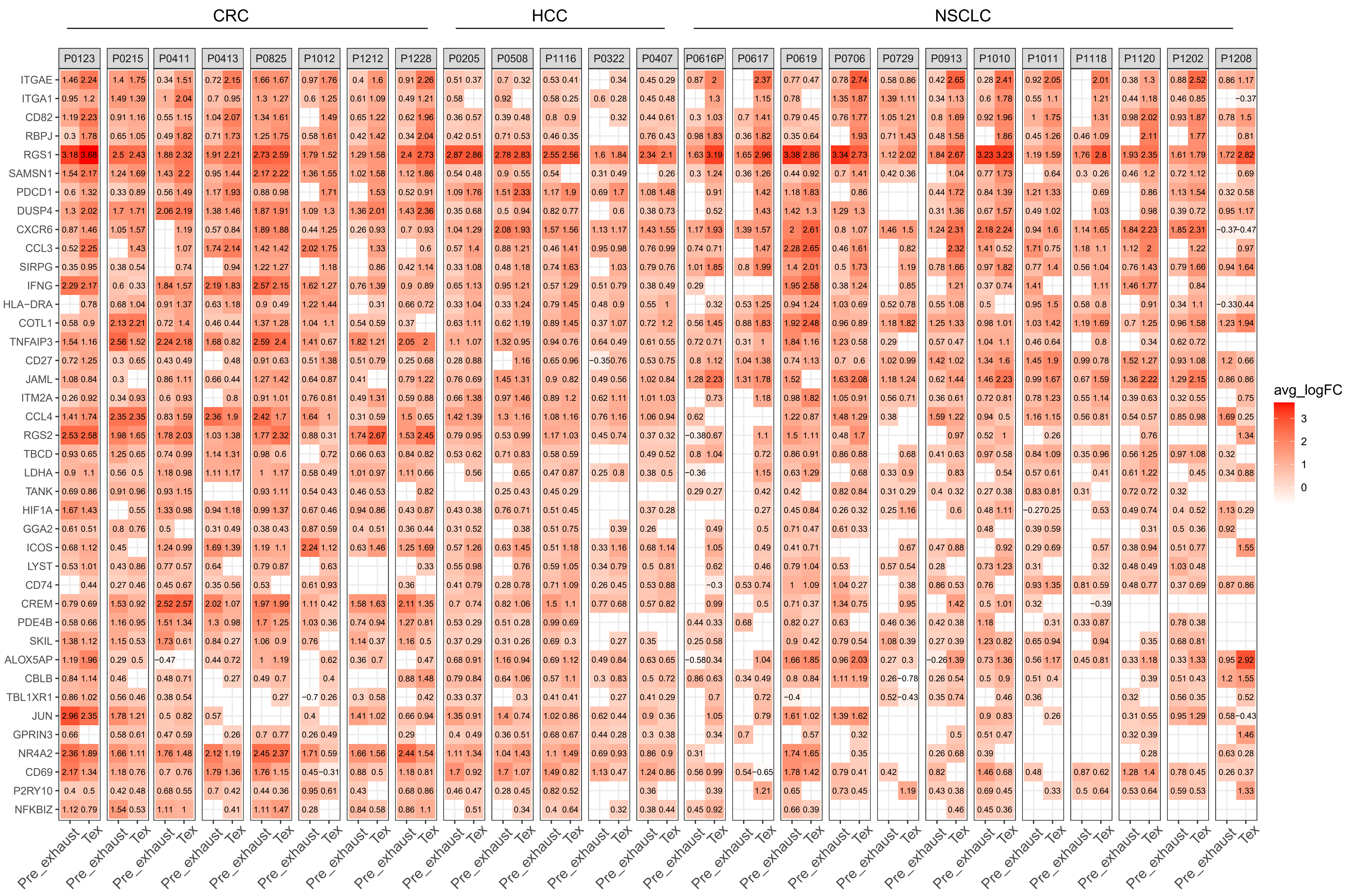

### Supplemental Figure 3

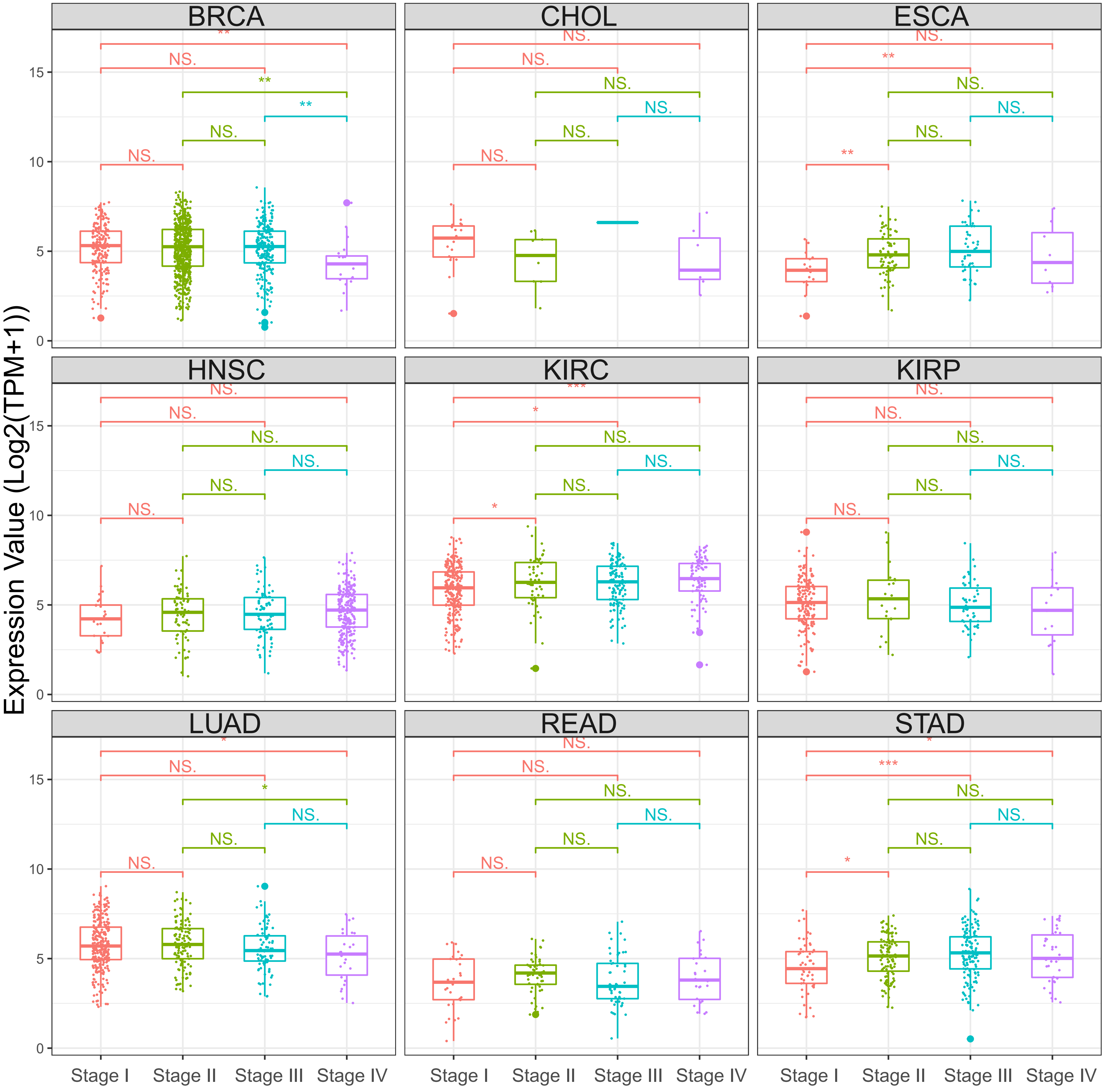

### Supplemental Figure 5

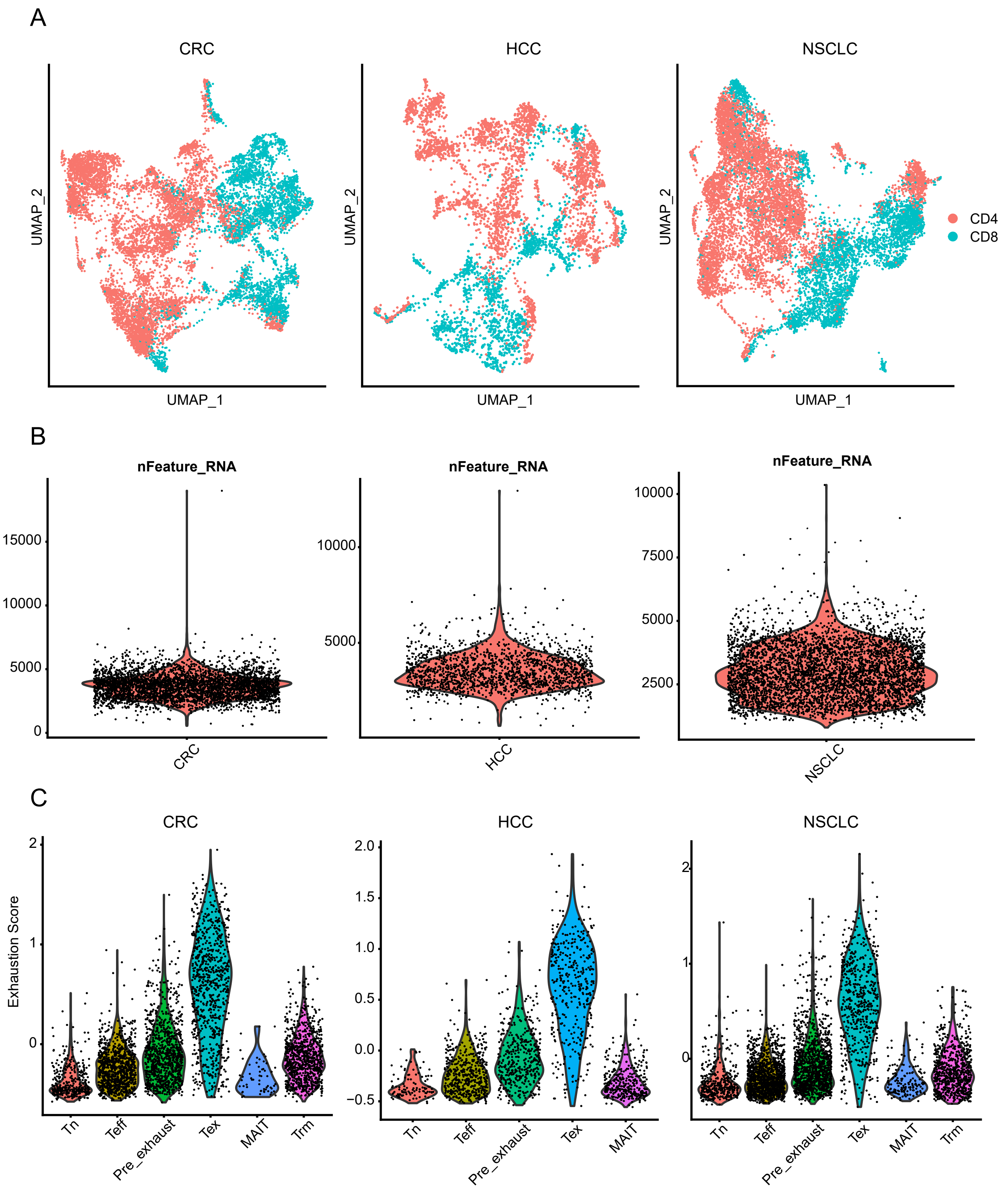
